# Social reward network connectivity differs between autistic and neurotypical youth during social interaction

**DOI:** 10.1101/2023.06.05.543807

**Authors:** Hua Xie, Dustin Moraczewski, Kathryn A. McNaughton, Katherine R. Warnell, Diana Alkire, Junaid S. Merchant, Laura A. Kirby, Heather A. Yarger, Elizabeth Redcay

## Abstract

A core feature of autism is difficulties with social interaction. Atypical social motivation is proposed to underlie these difficulties. However, prior work testing this hypothesis has shown mixed support and has been limited in its ability to understand real-world social-interactive processes in autism. We attempted to address these limitations by scanning neurotypical and autistic youth (n = 86) during a text-based reciprocal social interaction that mimics a “live” chat and elicits social reward processes. We focused on task-evoked functional connectivity (FC) of regions responsible for motivational-reward and mentalizing processes within the broader social reward circuitry. We found that task-evoked FC between these regions was significantly modulated by social interaction and receipt of social-interactive reward. Compared to neurotypical peers, autistic youth showed significantly greater task-evoked connectivity of core regions in the mentalizing network (e.g., posterior superior temporal sulcus) and the amygdala, a key node in the reward network. Furthermore, across groups, the connectivity strength between these mentalizing and reward regions was negatively correlated with self-reported social motivation and social reward during the scanner task. Our results highlight an important role of FC within the broader social reward circuitry for social-interactive reward. Specifically, greater context-dependent FC (i.e., differences between social engagement and non-social engagement) may indicate an increased “neural effort” during social reward and relate to differences in social motivation within autistic and neurotypical populations.

## 1. Introduction

Humans are inherently social creatures, relying on social interactions to survive and thrive, interactions that can be highly rewarding (Krach et al., 2010). Difficulty with social interactions is a central challenge and core diagnostic criterion for multiple psychiatric conditions, including autism (AUT, American Psychiatric Association, 2022). One controversial hypothesis is that these social challenges are due to differences in social motivation and social reward processing (Chevallier, Kohls, Troiani, Brodkin, & Schultz, 2012, but see Jaswal & Akhtar, 2018) and that these difficulties with social interaction may also be associated with an atypical neural circuitry (Clements et al., 2018). Thus, determining whether and how the neural substrates of social reward differ in typical and atypical development is critical to understanding the mechanisms underlying challenges in social interaction.

Numerous neuroimaging studies have examined the neural correlates of social reward processing, which involves the integration of regions within two networks (Ruff and Fehr, 2014). The first one is an overlapping cortical-basal-ganglia circuitry responsible for processing both monetary and social rewards (Huang et al., 2013). This shared motivational-reward network includes the striatum, orbitofrontal cortex, anterior cingulate cortex, and bilateral amygdala, supporting the idea of a “common neural currency” theory for reward processing (Montague and Berns, 2002; Wake and Izuma, 2017). Unlike monetary rewards, social reward processing in neurotypical (NT) participants also engages a broader social-cognitive network (Ruff and Fehr, 2014), including the temporoparietal junction, medial prefrontal cortex, superior temporal sulcus, and temporal pole, which we refer to here as the “mentalizing network.” The mentalizing network is associated with representing others’ mental states and is considered critical for efficient social interactions (Schurz et al., 2014). These two networks also tend to co-activate during social interaction (Alkire et al., 2018; Redcay and Schilbach, 2019; Xiao et al., 2022).

Despite theoretical arguments that atypical social reward circuitry may contribute to atypical social interactions in autism, empirical support is mixed. A recent meta-analysis surveying activation studies found that only a little more than half of the studies investigating social reward processing in autism (15 out of 27) showed atypical behavioral and/or physiological responses (Bottini, 2018).

Several factors may give rise to conflicting findings on the neural substrates of social reward processing in autism. One such factor is the heterogeneity among autistic (AUT) individuals^a^, including the level and display of social motivation (Wing, 1997). While autistic individuals often behave and express themselves in idiosyncratic ways, these behaviors do not necessarily indicate differences in motivation or desire for social connection (Jaswal and Akhtar, 2018). Some atypical behaviors interpreted as social disinterest could have alternative explanations unrelated to social motivation, such as anxiety and self-regulation difficulties (Kapp et al., 2011). Many autistic individuals also report high levels of loneliness and often long for friendship (Mazurek, 2014) – something that would be inconsistent with a “deficit” in social motivation. Thus, it is important to consider variability in subjective experiences of differences in social motivation and reward.

Another reason for prior mixed findings may be that the experimental paradigms used to study social reward often use non-interactive contexts. For example, a photo of a stranger’s smiling face is often used as a social reward. This lack of ecological validity is problematic (Redcay and Warnell, 2018) as recent studies have emphasized how participation in social interaction (as opposed to mere observation) alters the underlying cognitive and neural processing (Redcay and Schilbach, 2019; Schilbach et al., 2013). Therefore, researchers have increasingly advocated for assessing neural processes of social interaction by embedding the brain in a perceived live interactive setting (Redcay and Warnell, 2018), which may be critical to eliciting core social processing differences in autism (Rolison et al., 2015).

Relatedly, a potential methodological limitation of past work comes from their analytic approach. While past neuroimaging studies predominately focused on regional activation patterns to study social interaction and social reward, more recent studies have begun to investigate the brain’s functional coupling pattern using functional connectivity (FC), which is thought to reflect the brain’s interregional communication (van den Heuvel and Hulshoff Pol, 2010). Such studies have produced mixed findings on the neural signature of autism, with some reporting weaker connectivity in AUT compared to NT (“hypoconnectivity” in autism), others reporting greater connectivity in AUT (“hyperconnectivity”), and some claiming a combination of both (Hull et al., 2017). A few recent studies have begun to investigate how FC changes across cognitive states in AUT (e.g., Sridhar et al., 2021), termed as FC reconfiguration (for example, task-evoked conditions vs. null-task-demand conditions), as low FC reconfiguration may indicate that FC architecture easily adapts to task processing without significant ‘neural effort’ (Schultz & Cole, 2016). Moreover, while previous literature highlighted the critical role of the motivational-reward network and the mentalizing network for social reward processing, the interplay between the two has rarely been systematically investigated.

To address these gaps, we examined how the brain’s FC supports reward processing and social interaction, focusing on key brain regions within the social reward circuitry. Our experimental paradigm involved a “live” chat room, where neurotypical and autistic youth between middle childhood and early adolescence shared self-relevant information and received engaged responses from a peer or computer. This specific age range corresponds with considerable changes in youths’ social competencies and interpersonal relationships (Lam et al., 2014), offering a valuable window to understand underlying neural circuitry (Merchant et al., 2022). We hypothesized that connectivity patterns of the mentalizing and the motivational-reward network would be differentially modulated by social-interactive context and that the AUT group would show different connectivity patterns from the NT group. We further examined whether these differences can be accounted for by the heterogeneity of subjective experiences in social motivation and reward.

## 2 Method

### 2.1 Participants

Neurotypical participants were recruited using a database of families in the nearby metropolitan area and word of mouth. Autistic participants were recruited through local organizations, professional settings, listservs, and social media groups related to autism as well as word of mouth. Sixty-two autistic youth and ninety-nine neurotypical youth aged 7-14 years were recruited. The final sample (n = 86, see Appendix S1 for detailed inclusion criteria) included 43 autistic and 43 neurotypical youth, matched in age, gender, full-scale IQ, and in-scanner motion (Table 1). All procedures were approved by the Institutional Review Board of the University of Maryland, and parents and youth provided informed consent and assent.

**Table 1.** Matched sample characteristics (n = 86).

|  | NT (n=43) | AUT (n=43) | Statistics |
| --- | --- | --- | --- |
| Gender (#female/#male) | 8/35 | 7/36 | $\chi^2 (1, N = 86) = 0.09, p = 0.776$ |
| Age (yo, mean ± std) | 11.60 ± 1.90 | 11.86 ± 2.01 | $t(84) = -0.613, p = 0.542$ |
| Mean FD (mm) | 0.167 ± 0.068 | 0.179 ± 0.090 | $t(84) = -0.706, p = 0.482$ |
| Full scale IQ (FSIQ) | 116.6±14.4 | 113.4±14.9 | $t(84) = 1.010, p = 0.316$ |
| **Race** | NT (n=43) | AUT (n=43) | Total |
| Asian | 0 | 2 | 2 |
| Black or African American | 8 | 3 | 11 |
| White | 25 | 34 | 59 |
| More than One Race | 9 | 4 | 13 |
| Missing | 1 | 0 | 1 |
| **Ethnicity** |  |  |  |
| Hispanic/Latino | 5 | 1 | 6 |
| Not Hispanic/Latino | 36 | 42 | 78 |
| Missing/Does not wish to disclose | 2 | 0 | 2 |
| **Household Income** |  |  |  |
| \$75,000 or more | 37 | 38 | 75 |
| \$65,000-\$75,000 | 1 | 2 | 3 |
| \$45,000-\$65,000 | 1 | 2 | 3 |
| \$35,000-\$45,000 | 1 | 1 | 2 |
| Missing/Does not wish to disclose | 3 | 0 | 3 |

### 2.2 Experimental design

Youth participated in a real-time social interactive experiment designed to probe social reward systems (McNaughton et al., 2023; Warnell et al., 2018). As shown in Fig. 1, the participant engaged in a text-based ‘chat’ with an age- and gender-matched peer (who was actually simulated) by answering yes or no questions about themselves (e.g., “I play soccer”) followed by an engaged or disengaged response from the peer. They also completed computer trials, in which they shared information with the computer (for details, see Appendix S2).

**Figure. 1.**
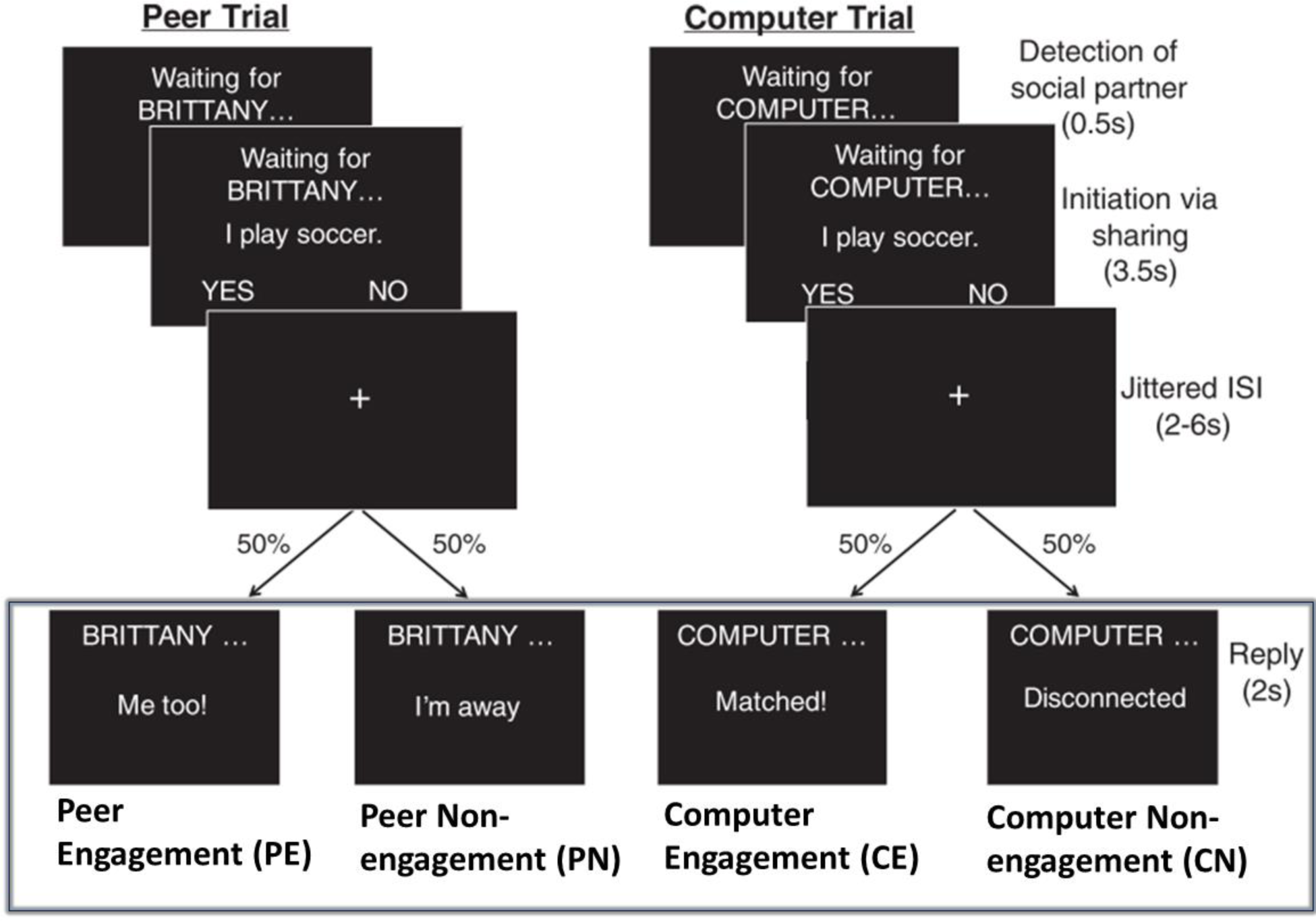
The interactive social interactive task with conditions of interest highlighted by the rectangle. Participants engaged in a text-based ‘chat’ with a peer or a computer by answering yes or no questions about themselves. The task consisted of two stages, initiation and reply. Here, we focused on the reply stage, and investigated three contrasts, i.e., social reward contrasts: PE vs. CE, PE vs. PN, and social context contrast: PE+PN vs. CE+CN (figure adapted from Warnell et al., (2018)).

Here, we focused on four types of reply events: Peer Engagement (PE), Peer Non-engagement (PN), Computer Engagement (CE), and Computer Non-engagement (CN), each with six trials per condition per run. The experiment was repeated over four runs, and each run lasted for approximately 6.2 mins. We included participants with three or more usable runs (n = 15 with 3 runs and n = 71 with 4 runs).

### 2.3 Post-scan interview

Immediately following the MRI scan, we conducted a verbal interview assessing how much the participants enjoyed interacting with the computer and peer on a 5-point Likert scale. Two post-scan enjoyment scores were used in this study, including a social motivation score (the difference between how much they wanted to see the answer from the peer vs. from the computer) and a social reward score (how much they liked chatting with the peer vs. computer; see Appendix S3 for details). We also assessed the participants’ belief in the live illusion by asking them if there was anything else they wanted to tell us. Seventeen participants who expressed disbelief were excluded from further analysis.

### 2.4 Image acquisition

The fMRI data were collected using a single Siemens 3T scanner 32-channel head coil at the Maryland Neuroimaging Center (MAGNETOM Trio Tim System, Siemens Medical Solutions). The scanning protocol consists of four functional runs (T2* weighted gradient echo-planar images; 40 interleaved axial slices; voxel size = 3.0 × 3.0 × 3.0 mm; repetition time = 2200 ms; echo time = 24 ms; flip angle = 78°; pixel matrix = 64 × 64), and one T1-weighted structural scan (176 sagittal slices, voxel size = 1.0 × 1.0 × 1.0 mm; repetition time = 1900 ms; echo time = 2.52 ms; flip angle = 9°; pixel matrix = 256 × 256).

### 2.5 fMRI data analysis

#### 2.5.1 Region of interests

Since our study primarily focuses on reward, we chose three a priori seed regions in the motivational-reward network, i.e., the bilateral amygdala, nucleus accumbens (NAcc), and ventral caudate (Fig. 2A). The amygdala was anatomically defined in Harvard-Oxford subcortical structural probability atlas in FSL, while NAcc and ventral caudate were described in a previous FC-based study (Di Martino et al., 2011). These regions were chosen given their contribution to social reward processing in autistic (Kohls et al., 2013b) and neurotypical youth (Ernst et al., 2005).

**Figure 2.**
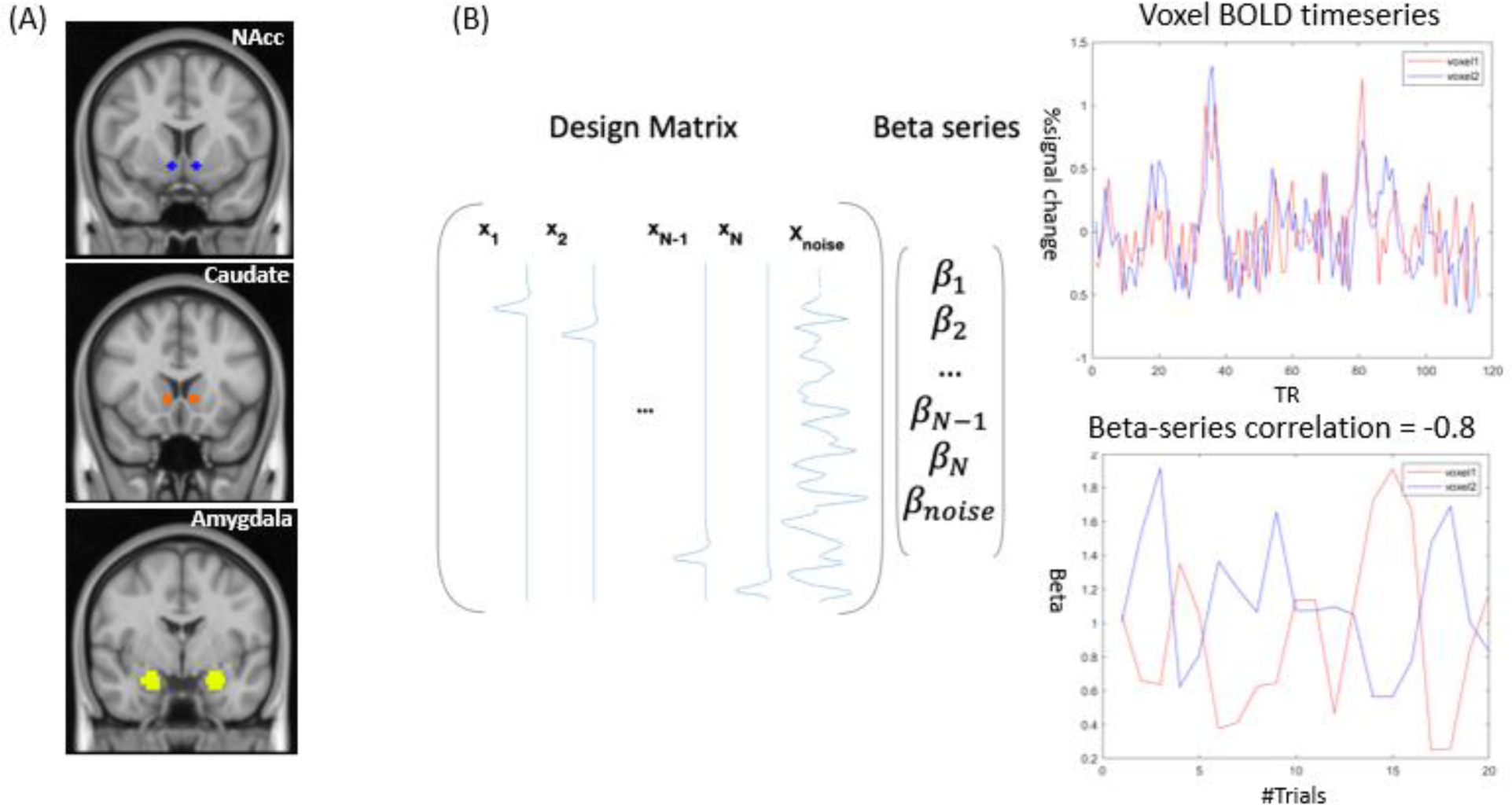
A graphical summary of the context-modulated FC analysis. (A) Three a priori seed regions are seed regions, i.e., nucleus accumbens (NAcc), ventral caudate, and amygdala. (B) Whole-brain context-modulated FC analysis. A new set of regressors were created to estimate trial-specific activation (beta coefficients). The context-modulated FC was estimated by correlating beta coefficients between the seed and voxels across the whole brain.

#### 2.5.2 Whole-brain context-modulated FC analysis

We evaluated the whole-brain context-modulated FC pattern using the subcortical ROIs as seeds with a beta-series connectivity approach. An overview of the analysis steps can be found in Fig. First, a separate regressor was created for each reply event, which was then convolved with a canonical HRF to model the BOLD response. Second, a trial-specific activation (beta coefficient) was estimated for each voxel using the Least Squares-All (LS-A) approach (Mumford et al., 2012), generating trial-wise regressors to identify trial-specific activation. We also censored the frames with FD greater than 1mm. Third, the beta series were averaged within each ROI, and the averaged timeseries were correlated with beta timeseries of all voxels using Spearman correlation to compute the whole-brain context-modulated FC. The correlation coefficients were subsequently Fisher-z transformed.

#### 2.5.3 General linear model analysis

To examine the effects of social context regardless of engagement, we examined the composite social context contrast by comparing the peer (PE+PN) to computer (CE+CN). To examine the effects of social reward specifically, we compared PE (i.e., peer engagement) — the social-interactive reward when participants receive an engaged, positive response from a peer — versus CE (computer engagement), where participants receive a response from a computer. Additionally, we compared PE vs. PN (i.e., peer non-engagement), where the peer is away and does not give a response. For simplicity, we only refer to the PE vs. CE contrast for the remainder of the paper when discussing our social reward contrast because we did not observe significant effects in PE vs. PN.

For each seed ROI, we conducted group-level analyses to identify voxels with significant main effects and group differences in the six whole-brain FC contrasts using AFNI function *3dMVM*, while controlling for age, gender, and the number of runs. To account for the multiple testing of three seeds and three contrasts, significant clusters were determined with a conservative cluster-wise false positive rate of 0.0056 (0.05/9 seeds, 124 voxels by *3dClustsim* based on average noise smoothness from the residual data) and a voxel-wise *p*-value of 0.001.

## 3. Results

### 3.1 Sample characteristics and behavioral findings

The characteristics of the matched samples are summarized in Table 1. A two-way analysis of variance test was used to examine the effects of interaction partner (peer vs. computer) and group (NT vs. AUT) on self-reported social reward and social motivation. We found a significant effect of social context such that both groups enjoyed interaction with peers more than the computer (*ps* < 0.001), but no effect of the group nor any interactions were found.

### 3.2 Context-modulated FC of social reward and social interaction

We used the trial-specific beta coefficients of three a priori seed regions. Below we report whole-brain context-modulated FC analyses for the social reward (i.e., social engagement vs. non-social engagement: PE vs. CE) and the social context contrast (i.e., chatting with a peer vs. computer: [PE + PN] vs. [CE + CN]). For the main effect of social context, we observed stronger connectivity between the left NAcc and the left inferior frontal gyrus (IFG) during social interaction (Fig. 3A). We also found significantly stronger FC in the AUT group between the amygdala seed and three regions: bilateral posterior superior temporal sulcus (pSTS) and right temporoparietal junction (TPJ) in the social reward contrast. All analyses were performed with a primary voxel-wise threshold of 0.001, and a cluster-wise threshold of 124 voxels, correcting for nine comparisons (3 seeds × 3 contrasts). No other significant effects survived cluster-wise correction. A summary of all significant clusters can be found in Table S1 in Supporting Information.

**Figure 3.**
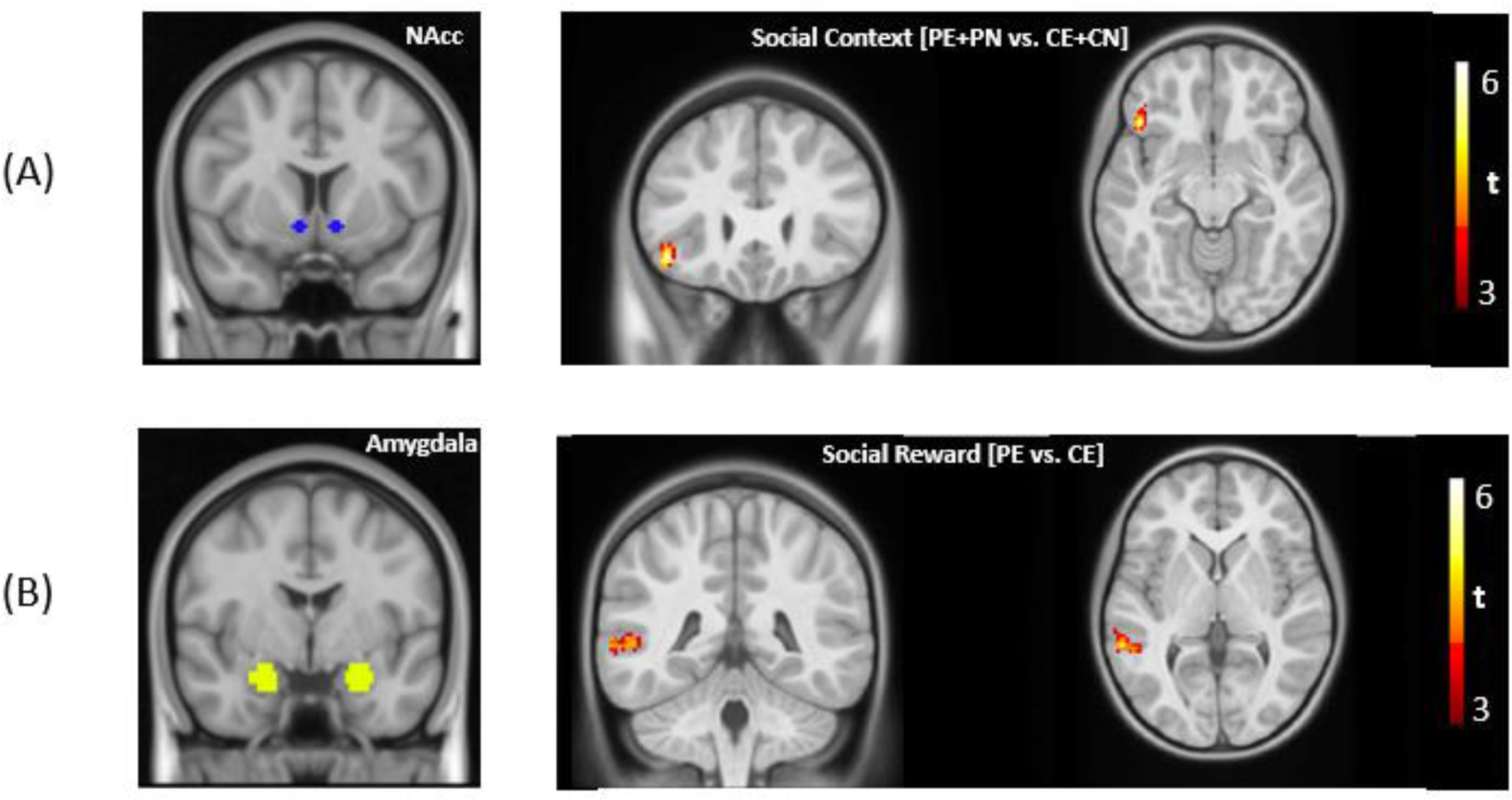
(A) For the main effects, significantly stronger FC was found between bilateral nucleus accumbens (NAcc) and left inferior frontal gyrus for the social context contrast. For the group differences, significantly stronger FC was found in the AUT group in the social reward contrast between the bilateral amygdala and bilateral pSTS and right TPJ, respectively (voxel-wise threshold = 0.001, cluster-wise threshold = 124 voxels).

### 3.3 Neural correlates of interaction enjoyment

To explore the mechanism underlying the significant group differences in context-modulated FC, we conducted post-hoc brain-behavior analyses between the FC values of the three significant clusters in the social reward contrast, and post-scan social reward and social motivation scores. As shown in Fig. 4A, after controlling for diagnosis, stronger FC between the amygdala and the left pSTS was related to lower social motivation scores (i.e., differences between how much participants wanted to see the answer from a peer vs. computer) within the combined sample (*t* = −2.24, *p* < 0.05). Similarly, as shown in Fig. 4B, FC between the amygdala and the right pSTS was negatively correlated with social reward scores (i.e., the differences between how much participants liked the answer from a peer or computer, *t* = −2.25, *p* < 0.05), after controlling for the diagnosis.

**Figure 4.**
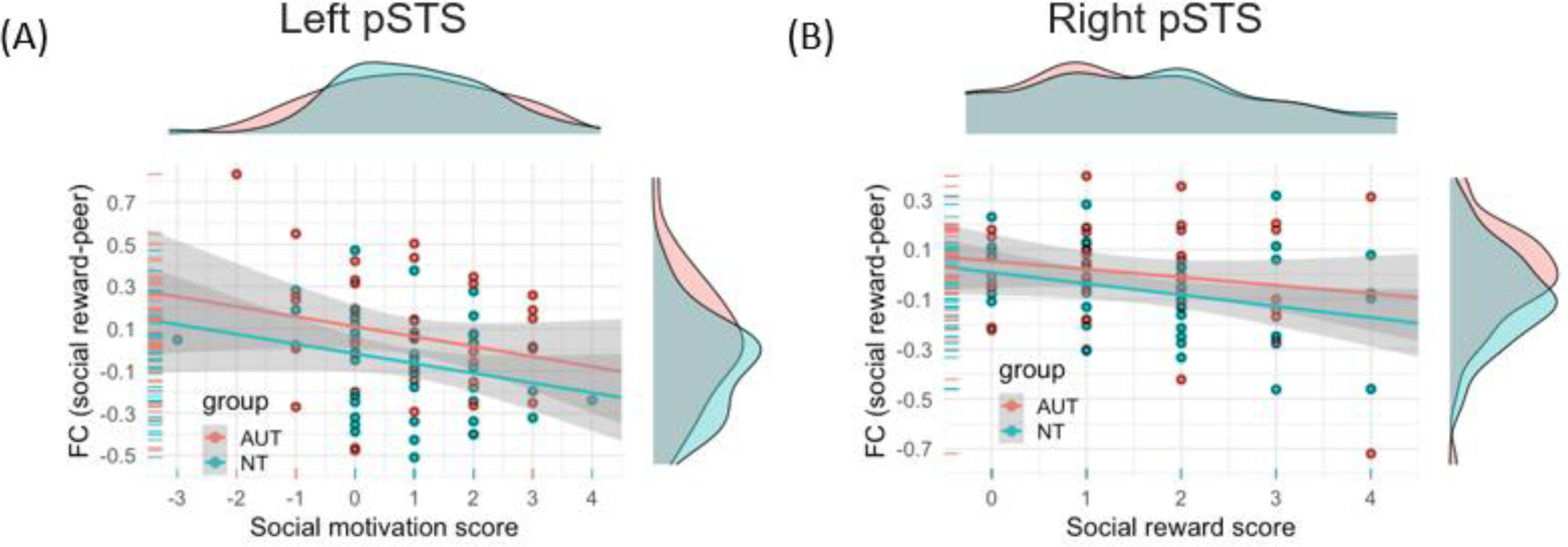
X-axis: Differences in post-scan enjoyment scores (peer – computer). Y-axis: FC of social reward contrasts (PE – CE). (A) A significant negative relationship was found between the differences in social motivation score and the FC between the amygdala and left pSTS (*t* = −2.24, *p* = 0.03). (B) A significant negative relationship was found between the differences in social reward score and the FC between the amygdala and right pSTS (*t* = −2.25, *p* = 0.03).

## Discussion

To understand whether social reward circuitry differs between autistic and neurotypical youth, we used a social-interactive paradigm to investigate task-induced FC changes during social reward processing in a sample of 86 youth between 7-14 years old. Middle childhood and early adolescence is a critical developmental period, as difficulties in social interaction during this period predict later mental health difficulties, poorer academic outcomes, and difficulties in later employment (Bornstein et al., 2010; Burt et al., 2008). Given earlier work highlighting the role of FC or co-activation between the mentalizing and reward networks during social interaction (Alkire et al., 2018; Assaf et al., 2013; Smith et al., 2014; Xiao et al., 2022), we hypothesized that connectivity profiles of regions associated with reward processing and mentalizing would be modulated by social context and that the AUT group would show different connectivity patterns in these regions. We found partial support for these hypotheses. Consistent with our hypotheses, we found greater connectivity within reward-relevant regions during social reward processing (i.e., receiving a positive response from a peer compared to a computer) in the full sample. We also found significant group differences in connectivity between key regions within the broader social reward circuitry (i.e., encompassing social-cognitive and reward-relevant regions of the amygdala and bilateral pSTS and R TPJ), and the amygdala-pSTS connectivity strength was related to individual differences in self-reported social motivation and reward across groups.

### Social context relates to enhanced mentalizing and reward network connectivity

For the main effect of social context (PE+PN vs. CE+CN, i.e., interacting with a peer vs. computer), consistent with our hypothesis, we found increased connectivity between a core component of the reward system (NAcc) and a part of the mentalizing network (left IFG). Left IFG is an important region associated with mentalizing and empathy (Arioli et al., 2021) and has also been found to encode youth’s interest levels in receiving feedback from peers (Guyer et al., 2012). Moreover, Pfeiffer and colleagues studied gaze-based interactions and found the activation in IFG and NAcc to be differentially modulated during interactions with a perceived human partner compared with a perceived computer (Pfeiffer et al., 2014). Therefore, a potential interpretation of this finding is that strengthened left IFG - NAcc connectivity is related to detecting and updating relevant social signals (particularly from interaction partners), with the NAcc encoding the reward prediction error (Schultz et al., 1997) and the IFG updating expectancies following social feedback (Kohls et al., 2013a; Nelson and Guyer, 2011). However, it is important to note that the IFG is a multi-functional region and this interpretation relies on a reverse inference (Poldrack, 2006).

### Stronger amygdala-mentalizing connectivity during peer engagement in AUT

We observed greater connectivity in the AUT group between the amygdala and bilateral pSTS and nearby right TPJ in the mentalizing network, when participants received a positive response from a peer compared to a computer (i.e., social reward contrast [PE vs. CE]). The amygdala is important for social cognition and social motivation (Chevallier et al., 2012; Kohls et al., 2013b), and the TPJ and pSTS are among key regions for social cognition and social interaction (Redcay et al., 2010). Our findings are consistent with prior neuroimaging studies demonstrating greater connectivity in autism (Chien et al., 2015; Dajani and Uddin, 2016; Fishman et al., 2014; Jasmin et al., 2019; Redcay et al., 2013; Supekar et al., 2013; Uddin et al., 2013; You et al., 2013), especially between subcortical and cortical regions (Cerliani et al., 2015; Ilioska et al., 2022).

More specifically, connectivity between pSTS/TPJ and the amygdala has often been highlighted in previous neuroimaging studies of reward-related processing in autism (Dichter, 2012). For example, Abrams and colleagues found weaker connectivity between human-voice-selective pSTS and the reward circuit, including the amygdala, in autistic children during rest (Abrams et al., 2013), indicating a potential role of amygdala-pSTS connectivity in reward and human voice processing. In contrast, stronger amygdala-STS connectivity was found in autism during spontaneous attention to eye gaze in emotional faces during a cognitive control task (Murphy et al., 2012), indicating the role of pSTS-amygdala connectivity in social information processing. Moreover, structural connectivity strength between the amygdala and pSTS was positively correlated with autistic traits (Iidaka et al., 2012).

Linking neural data to behavioral data may help us gain a more mechanistic understanding of the functional significance of the connectivity differences between groups (Picci et al., 2016). Interestingly, these group-level connectivity differences were present despite no group differences in behavioral self-report of either social motivation or social reward. Neural measures might prove to be more sensitive to identifying group differences than behavioral measures of social motivation/reward, due to potential confounds of bias in self-report (Van de Mortel, 2008) or observer expectations of social behavior (Jaswal and Akhtar, 2018). Importantly, we did find that behavioral reports tracked individual differences in connectivity, and this relation was similar in both groups. Specifically, connectivity between the amygdala and left and right pSTS was negatively correlated with self-reported social motivation and social reward scores. Thus, greater connectivity changes between conditions (i.e., greater FC reconfiguration) might reflect reduced social motivation or reduced sensitivity to social reward. We offer two possible interpretations for these observations: first, that greater FC reconfiguration indexes a domain-general difficulty switching between high-demanding and low-demanding conditions, and, second, that it may relate to social anxiety.

### Greater FC reconfiguration indexes greater neural effort?

First, greater FC reconfiguration between regions within the social reward circuitry may signal greater neural effort associated with social interaction (Schultz & Cole, 2016). Speaking to this possibility, a previous study found that high-demand social-interaction elicited greater connectivity between social brain regions in autistic compared to neurotypical youth in a live social interaction paradigm (Jasmin et al., 2019). When contrasting the condition with high social demand (conversation) with that of low social demand (repetition), they similarly found stronger task-evoked connectivity in social processing regions (e.g., STS) in autistic participants, and the increases in connectivity were positively related to the level of social impairment. Although these findings speak to differences in FC modulation in autism for social tasks, greater FC reconfiguration (i.e., task vs. control FC updates) has also been observed in autism during non-social cognitive tasks. For example, when contrasting a sustained attention task with rest, You et al. (2013) found that autistic children had increased distal connectivity between frontal, temporal, and parietal regions compared to neurotypical children, and this increased connectivity was associated with inattention problems in everyday life. In addition, Barttfeld et al. (2012) reported more pronounced connectivity changes in autistic adults than the neurotypical adults across three cognitive states with varying attention demands (i.e., rest, interoceptive, and exteroceptive attentional states). Lexical processing induces similarly greater FC configuration in autistic adolescents as compared with neurotypical peers (Sridhar et al., 2021). Beyond autism, Schultz & Cole (2016) found that neurotypical individuals with better task performance had smaller task-evoked FC reconfiguration when switching from task to rest, suggesting that better-performing individuals ‘pre-configure’ their FC at baseline (i.e., rest) to be more efficiently updated for various processing demands. Taken together, greater FC reconfiguration may indicate that social interaction requires greater neural effort and thus is mentally taxing for individuals who find social interaction difficult, regardless of diagnostic status.

### Strengthened amygdala connectivity reflects social anxiety?

A second potential explanation for stronger connectivity between the amygdala and mentalizing regions relates to social anxiety. The amygdala is a multi-functional brain region. While here we chose the amygdala as a seed given its central role in reward processing and social motivation, the amygdala is also known for its involvement in regulating emotions such as anxiety (Davidson, 2002). Moreover, anxiety disorders often co-occur with autism, as evidenced by a recent meta-analysis showing about 40% of autistic youths have received a diagnosis of at least one anxiety disorder (van Steensel et al., 2011), while atypical amygdala connectivity is also common in these anxiety disorders (Sylvester et al., 2012). For example, in adults with social anxiety disorder, Pannekoek and colleagues found heightened resting-state connectivity between the right amygdala and the left middle temporal gyrus overlapping with our left pSTS cluster (Pannekoek et al., 2013). Thus, it is possible that for autistic youth, especially those with social anxiety or a history of peer rejection, the social stimulus could be more anxiety-inducing, causing heightened amygdala connectivity. To test this possibility, we reran the analysis after excluding the eight autistic youth with anxiety disorders. We found similar significant group differences in right pSTS and TPJ as well as significant brain-behavior relationships with interaction enjoyment, which does not offer direct support for this possibility. Furthermore, our post-hoc exploratory analysis using the parent-reported social phobia scale of the Screen for Child Anxiety Related Emotional Disorders (Birmaher et al., 1997) failed to find any significant relationship with connectivity between the amygdala and mentalizing regions within a subset of participants (n = 56). However, given the limited sample size, the case-control study design, and the prevalence of subclinical anxiety in youth (Vasa et al., 2013), we encourage further studies with larger samples of participants with anxiety (with and without autism) and investigation of whether social anxiety traits are driving the relation we found between mentalizing-reward network connectivity and social reward and motivation.

### Limitations and future directions

The current study focused on individual differences in processing interactive social reward, including self-report measures of social motivation and reward. However, the differences in social abilities for autistic youth may also stem from other factors, such as theory of mind, executive function, or sensory processing differences. Moreover, the interactive chat task used in the current study is text-based, and text-based communication may alleviate some of the difficulties autistic people experience in face-to-face contexts (Benford and Standen, 2009). The task also is highly structured, which significantly reduces uncertainty (Boulter et al., 2014). Thus, our highly structured, text-based interaction may have diminished some potential group differences. Additionally, given the exploratory nature of our post-hoc brain-behavioral analysis, we did not perform any multiple comparisons correction, and future work is needed to validate our findings. Lastly, although we controlled for age in our analysis, age may add more variance, as previous studies have suggested that autistic youth from different age cohorts may show different connectivity patterns (Uddin et al., 2013).

## Conclusion

In sum, our study demonstrated increased integration between regions associated with reward and mentalizing networks during social reward processing in a live reciprocal social interaction. The greater task-based modulation of connectivity (i.e., greater reconfiguration) within the broader social reward circuitry may contribute to group and individual differences in how social-interactive reward is processed.

## Acknowledgments

Research reported in this publication was supported by the National Institute of Mental Health of the National Institutes of Health under award number R01MH107441. We thank the participants and their families, Maryland Neuroimaging Center and staff for support with data collection, and members of our research team who assisted with data collection, including Kayla Velnoskey, Sydney Maniscalco, Aiste Cechaviciute, Jacqueline Thomas, Alexandra Hickey, Eleonora Sadikova, and Meredith Pecukonis.

## Supplemental Materials

### S1. Inclusion criteria

Youth were excluded if they were born premature (<34 weeks), non-native English speakers, had a history of concussion or head trauma, or had a full-scale IQ < 80 as assessed by the Kaufman Brief Intelligence Test, Second Edition (Kaufman, 2004). Parents of neurotypical youth reported no history of neurological or psychiatric disorders or first-degree relatives with autism or schizophrenia. Autistic youth were not excluded if parents reported a common co-occurring mental health condition, including attention-deficit/hyperactivity disorder (n = 19), obsessive-compulsive disorder (n = 1), or anxiety (n = 8). Autistic youth were eligible to participate only if they had a prior clinical diagnosis of autism which was then confirmed by our research team using the Autism Diagnostic Observation Schedule, 2nd edition (ADOS-2, Lord *et al*., 2012) via a licensed clinical psychologist or clinical psychology graduate student who was research-reliable in ADOS administration and coding. Of youth who completed the MRI scan, the following inclusion criteria were used: believed they were chatting with a real peer partner (see section 2.3 Post-scan interview), and adequate task performance (i.e., responding to at least 2/3 of trials) and with three or more usable runs (i.e., mean framewise displacement (FD) < 0.5mm and maximum FD < 5.2mm, corresponding to the diagonal length of a 3mm isomorphic cube). The final sample (n = 86) overlaps with the sample used in McNaughton *et al*. (2023).

### S2. Experiment

Participants were first informed whether the recipient was a peer (peer trial) or a computer (computer trial), both of which had an equal possibility. Then, the participants initiated an interaction by answering a Yes/No question about their likes and hobbies (e.g., “I like soccer”). After a jittering 2-6 sec (mean 3.5 sec) fixation period, participants received a response in the two-second reply phase. The responses consisted of engagement (e.g., “Me too!”, indicating the peer agreed with the participant, or “Matched!”, indicating that the computer randomly generated the same answer as the child), non-engagement (e.g., “I’m away” or “Disconnected”), and disagreement (e.g., “That’s not what I picked”).

For peer trials, the participants were told that the peer would sometimes be unable to respond because the peer was playing another game. For these disengaged trials, an away message was displayed as the peer response. Moreover, participants believed that the peer always saw their answer, and the peer would either respond if they were able to or not respond if they had been assigned to play another game. For computer trials, participants believed that the computer would randomly pick an answer following participants’ answering the Yes/No question. Moreover, participants were informed that the computer would sometimes lose the connection and be unable to generate an answer, resulting in disengage trials. The order and timing of trials were predetermined, and we used four sets of stimuli to avoid pairing the questions with reply types (e.g., the “I play soccer” trials did not always receive a “Me too” response).

### S3. Post-scan enjoyment questionnaire

A post-scan questionnaire was filled by participants using a 1-5 Likert scale (1 = not at all, to 5 = a lot). The items used in the analysis are listed below.

How much did you want to see his/her answer to your question?

How much did you want to see if the computer matched your answer?

How much did you like chatting with ?

How much did you like it when you were just answering the computer?

### S4. Preprocessing

A standardized preprocessing pipeline fMRIprep v1.4.1 was used to preprocess the imaging data (Esteban *et al*., 2019). The skull-stripped BOLD images underwent motion correction, slice timing correction, and susceptibility distortion correction, and were lastly resampled to MNI space. Automatic removal of motion artifacts using independent component analysis was performed on the preprocessed BOLD images after removal of non-steady-state volumes and spatial smoothing with an isotropic, Gaussian kernel of 6mm FWHM (full-width half-maximum). Lastly, the BOLD images were then intensity normalized to have a mean intensity of 1,000, and a binary group mask at the threshold of 0.9 probability was applied.

**Table S1.**
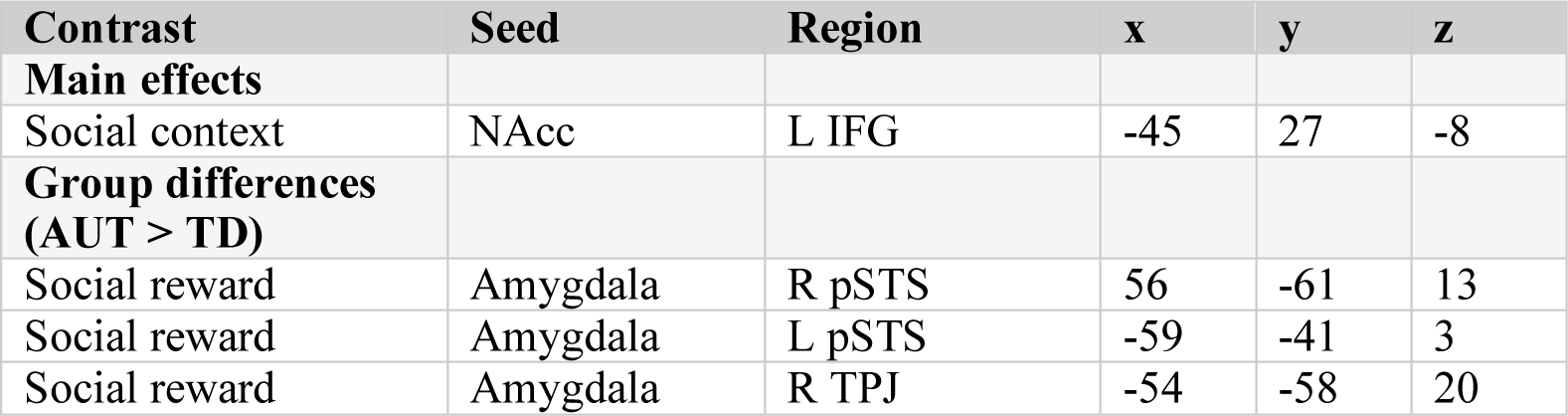
Clusters with significant main effect and group differences with MNI coordinates of their center of mass (voxel-wise p-value = 0.001, cluster-wise threshold = 124 voxels).

| Contrast | Seed | Region | x | y | z |
| --- | --- | --- | --- | --- | --- |
| **Main effects** |  |  |  |  |  |
| Social context | NAcc | L IFG | -45 | 27 | -8 |
| **Group differences (AUT > TD)** |  |  |  |  |  |
| Social reward | Amygdala | R pSTS | 56 | -61 | 13 |
| Social reward | Amygdala | L pSTS | -59 | -41 | 3 |
| Social reward | Amygdala | R TPJ | -54 | -58 | 20 |

## Footnotes

a We acknowledge preferences for different languages used when referring to autism. Due to a reported preference for identity-first language among a majority of autistic individuals and caregivers, we use that in this article.

## Reference

Abrams, D.A., Lynch, C.J., Cheng, K.M., Phillips, J., Supekar, K., Ryali, S., Uddin, L.Q., Menon, V., 2013. Underconnectivity between voice-selective cortex and reward circuitry in children with autism. Proc. Natl. Acad. Sci. U. S. A. 110, 12060–12065. https://doi.org/10.1073/pnas.1302982110

Alkire, D., Levitas, D., Warnell, K.R., Redcay, E., 2018. Social interaction recruits mentalizing and reward systems in middle childhood. Hum. Brain Mapp. 39, 3928–3942. https://doi.org/10.1002/hbm.24221

American Psychiatric Association, 2022. Diagnostic and Statistical Manual of Mental Disorders: DSM-5-TR (Fifth edition, text revision), The Curated Reference Collection in Neuroscience and Biobehavioral Psychology. American Psychiatric Pub.

Arioli, M., Cattaneo, Z., Ricciardi, E., Canessa, N., 2021. Overlapping and specific neural correlates for empathizing, affective mentalizing, and cognitive mentalizing: A coordinate-based meta-analytic study. Hum. Brain Mapp. 42, 4777–4804. https://doi.org/10.1002/hbm.25570

Assaf, M., Hyatt, C.J., Wong, C.G., Johnson, M.R., Schultz, R.T., Hendler, T., Pearlson, G.D., 2013. Mentalizing and motivation neural function during social interactions in autism spectrum disorders. NeuroImage Clin. 3, 321–331. https://doi.org/10.1016/j.nicl.2013.09.005

Barttfeld, P., Wicker, B., Cukier, S., Navarta, S., Lew, S., Leiguarda, R., Sigman, M., 2012. State-dependent changes of connectivity patterns and functional brain network topology in autism spectrum disorder. Neuropsychologia 50, 3653–3662. https://doi.org/10.1016/j.neuropsychologia.2012.09.047

Benford, P., Standen, P., 2009. The internet: A comfortable communication medium for people with Asperger syndrome (AS) and high functioning autism (HFA)? J. Assist. Technol. 3, 44–53. https://doi.org/10.1108/17549450200900015

Birmaher, B., Khetarpal, S., Brent, D., Cully, M., Balach, L., Kaufman, J., Neer, S.M., 1997. The screen for child anxiety related emotional disorders (SCARED): Scale construction and psychometric characteristics. J. Am. Acad. Child Adolesc. Psychiatry 36, 545–553.

Bornstein, M.H., Hahn, C.S., Haynes, O.M., 2010. Social competence, externalizing, and internalizing behavioral adjustment from early childhood through early adolescence: Developmental cascades. Dev. Psychopathol. 22, 717–735. https://doi.org/10.1017/S0954579410000416

Bottini, S., 2018. Social reward processing in individuals with autism spectrum disorder: A systematic review of the social motivation hypothesis. Res. Autism Spectr. Disord. 45, 9–26. https://doi.org/10.1016/j.rasd.2017.10.001

Boulter, C., Freeston, M., South, M., Rodgers, J., 2014. Intolerance of uncertainty as a framework for understanding anxiety in children and adolescents with autism spectrum disorders. J. Autism Dev. Disord. 44, 1391–1402.

Burt, K.B., Obradović, J., Long, J.D., Masten, A.S., 2008. The interplay of social competence and psychopathology over 20 years: Testing transactional and cascade models. Child Dev. 79, 359–374.

Cerliani, L., Mennes, M., Thomas, R.M., Di Martino, A., Thioux, M., Keysers, C., 2015. Increased functional connectivity between subcortical and cortical resting-state networks in Autism spectrum disorder. JAMA Psychiatry 72, 767–777. https://doi.org/10.1001/jamapsychiatry.2015.0101

Chevallier, C., Kohls, G., Troiani, V., Brodkin, E.S., Schultz, R.T., 2012. The social motivation theory of autism. Trends Cogn. Sci. 16, 231–239. https://doi.org/10.1016/j.tics.2012.02.007

Chien, H., Lin, H., Lai, M., Gau, S.S., Tseng, W.I., 2015. Hyperconnectivity of the right posterior temporo-parietal junction predicts social difficulties in boys with autism spectrum disorder. Autism Res. 8, 427–441.

Clements, C.C., Zoltowski, A.R., Yankowitz, L.D., Yerys, B.E., Schultz, R.T., Herrington, J.D., 2018. Evaluation of the social motivation hypothesis of autism a systematic review and meta-analysis. JAMA Psychiatry 75, 797–808. https://doi.org/10.1001/jamapsychiatry.2018.1100

Dajani, D.R., Uddin, L.Q., 2016. Local brain connectivity across development in autism spectrum disorder: A cross-sectional investigation. Autism Res. 9, 43–54. https://doi.org/10.1002/aur.1494

Davidson, R.J., 2002. Anxiety and affective style: Role of prefrontal cortex and amygdala. Biol. Psychiatry 51, 68–80. https://doi.org/10.1016/S0006-3223(01)01328-2

Di Martino, A., Kelly, C., Grzadzinski, R., Zuo, X.N., Mennes, M., Mairena, M.A., Lord, C., Castellanos, F.X., Milham, M.P., 2011. Aberrant striatal functional connectivity in children with autism. Biol. Psychiatry 69, 847–856. https://doi.org/10.1016/j.biopsych.2010.10.029

Dichter, G.S., 2012. Functional magnetic resonance imaging of autism spectrum disorders. Dialogues Clin. Neurosci. 14, 319–351. https://doi.org/10.31887/dcns.2012.14.3/gdichter

Ernst, M., Nelson, E.E., Jazbec, S., McClure, E.B., Monk, C.S., Leibenluft, E., Blair, J., Pine, D.S., 2005. Amygdala and nucleus accumbens in responses to receipt and omission of gains in adults and adolescents. Neuroimage 25, 1279–1291. https://doi.org/10.1016/j.neuroimage.2004.12.038

Fishman, I., Keown, C.L., Lincoln, A.J., Pineda, J.A., Müller, R.A., 2014. Atypical cross talk between mentalizing and mirror neuron networks in autism spectrum disorder. JAMA Psychiatry 71, 751– 760. https://doi.org/10.1001/jamapsychiatry.2014.83

Guyer, A.E., Choate, V.R., Pine, D.S., Nelson, E.E., 2012. Neural circuitry underlying affective response to peer feedback in adolescence. Soc. Cogn. Affect. Neurosci. 7, 81–92. https://doi.org/10.1093/scan/nsr043

Huang, P., Qiu, L., Shen, L., Zhang, Y., Song, Z., Qi, Z., Gong, Q., Xie, P., 2013. Evidence for a left-over-right inhibitory mechanism during figural creative thinking in healthy nonartists. Hum. Brain Mapp. 34, 2724–2732. https://doi.org/10.1002/hbm.22093

Hull, J. V., Jacokes, Z.J., Torgerson, C.M., Irimia, A., Van Horn, J.D., Aylward, E., Bernier, R., Bookheimer, S., Dapretto, M., Gaab, N., Geschwind, D., Jack, A., Nelson, C., Pelphrey, K., State, M., Ventola, P., Webb, S.J., 2017. Resting-state functional connectivity in autism spectrum disorders: A review. Front. Psychiatry 7. https://doi.org/10.3389/fpsyt.2016.00205

Iidaka, T., Miyakoshi, M., Harada, T., Nakai, T., 2012. White matter connectivity between superior temporal sulcus and amygdala is associated with autistic trait in healthy humans. Neurosci. Lett. 510, 154–158. https://doi.org/10.1016/j.neulet.2012.01.029

Ilioska, I., Oldehinkel, M., Llera, A., Chopra, S., Looden, T., Chauvin, R., Rooij, D. Van, Floris, D.L., Tillmann, J., Moessnang, C., Banaschewski, T., Holt, R.J., Loth, E., Charman, T., Murphy, D.G.M., Ecker, C., Mennes, M., Beckmann, C.F., Fornito, A., Buitelaar, J.K., group, the E.-A.L., 2022. Connectome-wide mega-analysis reveals robust patterns of atypical functional connectivity in autism. medRxiv 2022.01.09.22268936.

Jasmin, K., Gotts, S.J., Xu, Y., Liu, S., Riddell, C.D., Ingeholm, J.E., Kenworthy, L., Wallace, G.L., Braun, A.R., Martin, A., 2019. Overt social interaction and resting state in young adult males with autism: Core and contextual neural features. Brain 142, 808–822. https://doi.org/10.1093/brain/awz003

Jaswal, V.K., Akhtar, N., 2018. Being versus appearing socially uninterested: Challenging assumptions about social motivation in autism. Behav. Brain Sci. 42, 1–84. https://doi.org/10.1017/S0140525X18001826

Kapp, S.K., Gantman, A., Laugeson, E.A., 2011. Transition to adulthood for high-functioning individuals with autism spectrum disorders, A comprehensive book on autism spectrum disorders. InTech Rijeka. https://doi.org/10.5772/21506

Kohls, G., Perino, M.T., Taylor, J.M., Madva, E.N., Cayless, S.J., Troiani, V., Price, E., Faja, S., Herrington, J.D., Schultz, R.T., 2013a. The nucleus accumbens is involved in both the pursuit of social reward and the avoidance of social punishment. Neuropsychologia 51, 2062–2069. https://doi.org/10.1016/j.neuropsychologia.2013.07.020

Kohls, G., Schulte-Rüther, M., Nehrkorn, B., Müller, K., Fink, G.R., Kamp-Becker, I., Herpertz-Dahlmann, B., Schultz, R.T., Konrad, K., 2013b. Reward system dysfunction in autism spectrum disorders. Soc. Cogn. Affect. Neurosci. 8, 565–572. https://doi.org/10.1093/scan/nss033

Krach, S., Paulus, F.M., Bodden, M., Kircher, T., 2010. The rewarding nature of social interactions. Front. Behav. Neurosci. 4. https://doi.org/10.3389/fnbeh.2010.00022

Lam, C.B., Mchale, S.M., Crouter, A.C., 2014. Time with peers from middle childhood to late adolescence: Developmental course and adjustment correlates. Child Dev. 85, 1677–1693. https://doi.org/10.1111/cdev.12235

Mazurek, M.O., 2014. Loneliness, friendship, and well-being in adults with autism spectrum disorders. Autism 18, 223–232. https://doi.org/10.1177/1362361312474121

McNaughton, K.A., Kirby, L.A., Warnell, K.R., Alkire, D., Merchant, J.S., Moraczewski, D., Yarger, H.A., Thurm, A., Redcay, E., 2023. Social-interactive reward elicits similar neural response in autism and typical development and predicts future social experiences. Dev. Cogn. Neurosci. 59, 101197. https://doi.org/10.1016/j.dcn.2023.101197

Merchant, J.S., Alkire, D., Redcay, E., 2022. Neural similarity between mentalizing and live social interaction during the transition to adolescence. Hum. Brain Mapp. 1–17. https://doi.org/10.1002/hbm.25903

Montague, P.R., Berns, G.S., 2002. Neural economics and the biological substrates of valuation. Neuron 36, 265–284. https://doi.org/10.1016/S0896-6273(02)00974-1

Mumford, J.A., Turner, B.O., Ashby, F.G., Poldrack, R.A., 2012. Deconvolving BOLD activation in event-related designs for multivoxel pattern classification analyses. Neuroimage 59, 2636–2643. https://doi.org/10.1016/j.neuroimage.2011.08.076

Murphy, E.R., Foss-Feig, J., Kenworthy, L., Gaillard, W.D., Vaidya, C.J., 2012. Atypical Functional Connectivity of the Amygdala in Childhood Autism Spectrum Disorders during Spontaneous Attention to Eye-Gaze. Autism Res. Treat. 2012, 1–12. https://doi.org/10.1155/2012/652408

Nelson, E.E., Guyer, A.E., 2011. The development of the ventral prefrontal cortex and social flexibility. Dev. Cogn. Neurosci. 1, 233–245. https://doi.org/10.1016/j.dcn.2011.01.002

Pannekoek, J.N., Veer, I.M., Van Tol, M.J., Van der Werff, S.J.A., Demenescu, L.R., Aleman, A., Veltman, D.J., Zitman, F.G., Rombouts, S.A.R.B., Van der Wee, N.J.A., 2013. Resting-state functional connectivity abnormalities in limbic and salience networks in social anxiety disorder without comorbidity. Eur. Neuropsychopharmacol. 23, 186–195. https://doi.org/10.1016/j.euroneuro.2012.04.018

Pfeiffer, U.J., Schilbach, L., Timmermans, B., Kuzmanovic, B., Georgescu, A.L., Bente, G., Vogeley, K., 2014. Why we interact: On the functional role of the striatum in the subjective experience of social interaction. Neuroimage 101, 124–137. https://doi.org/10.1016/j.neuroimage.2014.06.061

Picci, G., Gotts, S.J., Scherf, K.S., 2016. A theoretical rut: revisiting and critically evaluating the generalized under/over-connectivity hypothesis of autism. Dev. Sci. 19, 524–549. https://doi.org/10.1111/desc.12467

Poldrack, R.A., 2006. Can cognitive processes be inferred from neuroimaging data? Trends Cogn. Sci. 10, 59–63. https://doi.org/10.1016/j.tics.2005.12.004

Redcay, E., Dodell-Feder, D., Pearrow, M.J., Mavros, P.L., Kleiner, M., Gabrieli, J.D.E., Saxe, R., 2010. Live face-to-face interaction during fMRI: A new tool for social cognitive neuroscience. Neuroimage 50, 1639–1647. https://doi.org/10.1016/j.neuroimage.2010.01.052

Redcay, E., Moran, J.M., Mavros, P.L., Tager-Flusberg, H., Gabrieli, J.D.E., Whitfield-Gabrieli, S., 2013. Intrinsic functional network organization in high-functioning adolescents with autism spectrum disorder. Front. Hum. Neurosci. 7, 1–11. https://doi.org/10.3389/fnhum.2013.00573

Redcay, E., Schilbach, L., 2019. Using second-person neuroscience to elucidate the mechanisms of social interaction. Nat. Rev. Neurosci. 20, 495–505. https://doi.org/10.1038/s41583-019-0179-4

Redcay, E., Warnell, K.R., 2018. A Social-Interactive Neuroscience Approach to Understanding the Developing Brain, 1st ed, Advances in Child Development and Behavior. Elsevier Inc. https://doi.org/10.1016/bs.acdb.2017.10.001

Rolison, M.J., Naples, A.J., McPartland, J.C., 2015. Interactive social neuroscience to study autism spectrum disorder. Yale J. Biol. Med. 88, 17–24.

Ruff, C.C., Fehr, E., 2014. The neurobiology of rewards and values in social decision making. Nat. Rev. Neurosci. 15, 549–562. https://doi.org/10.1038/nrn3776

Schilbach, L., Timmermans, B., Reddy, V., Costall, A., Bente, G., Schlicht, T., Vogeley, K., 2013. Toward a second-person neuroscience. Behav. Brain Sci. 36, 393–414. https://doi.org/10.1017/S0140525X12000660

Schultz, D.H., Cole, M.W., 2016. Higher Intelligence Is Associated with Less Task-Related Brain Network Reconfiguration. J. Neurosci. 36, 8551–8561. https://doi.org/10.1523/JNEUROSCI.0358-16.2016

Schultz, W., Dayan, P., Montague, P.R., 1997. A neural substrate of prediction and reward. Science (80-.). 275, 1593–1599. https://doi.org/10.1126/science.275.5306.1593

Schurz, M., Radua, J., Aichhorn, M., Richlan, F., Perner, J., 2014. Fractionating theory of mind: A meta-analysis of functional brain imaging studies. Neurosci. Biobehav. Rev. https://doi.org/10.1016/j.neubiorev.2014.01.009

Smith, D. V., Clithero, J.A., Boltuck, S.E., Huettel, S.A., 2014. Functional connectivity with ventromedial prefrontal cortex reflects subjective value for social rewards. Soc. Cogn. Affect. Neurosci. 9, 2017–2025. https://doi.org/10.1093/scan/nsu005

Sridhar, A., Keehn, R., Jao, J., Wilkinson, M., Gao, Y., Olson, M., Mash, L.E., Alemu, K., Manley, A., Marinkovic, K., Linke, A., Müller, R.-A., 2021. Increased heterogeneity and task-related reconfiguration of functional connectivity within a lexicosemantic network in autism 1–50.

Supekar, K., Uddin, L.Q., Khouzam, A., Phillips, J., Gaillard, W.D., Kenworthy, L.E., Yerys, B.E., Vaidya, C.J., Menon, V., 2013. Brain Hyperconnectivity in Children with Autism and its Links to Social Deficits. Cell Rep. 5, 738–747. https://doi.org/10.1016/j.celrep.2013.10.001

Sylvester, C.M., Corbetta, M., Raichle, M.E., Rodebaugh, T.L., Schlaggar, B.L., Sheline, Y.I., Zorumski, C.F., Lenze, E.J., 2012. Functional network dysfunction in anxiety and anxiety disorders. Trends Neurosci. 35, 527–535. https://doi.org/10.1016/j.tins.2012.04.012

Uddin, L.Q., Supekar, K., Menon, V., 2013. Reconceptualizing functional brain connectivity in autism from a developmental perspective. Front. Hum. Neurosci. 7, 1–11. https://doi.org/10.3389/fnhum.2013.00458

Van de Mortel, T.F., 2008. Faking it: social desirability response bias in self-report research. Aust. J. Adv. Nursing, 25, 40–48.

van den Heuvel, M.P., Hulshoff Pol, H.E., 2010. Exploring the brain network: A review on resting-state fMRI functional connectivity. Eur. Neuropsychopharmacol. 20, 519–534. https://doi.org/10.1016/j.euroneuro.2010.03.008

van Steensel, F.J.A., Bögels, S.M., Perrin, S., 2011. Anxiety Disorders in Children and Adolescents with Autistic Spectrum Disorders: A Meta-Analysis. Clin. Child Fam. Psychol. Rev. 14, 302–317. https://doi.org/10.1007/s10567-011-0097-0

Vasa, R.A., Kalb, L., Mazurek, M., Kanne, S., Freedman, B., Keefer, A., Clemons, T., Murray, D., 2013. Age-related differences in the prevalence and correlates of anxiety in youth with autism spectrum disorders. Res. Autism Spectr. Disord. 7, 1358–1369. https://doi.org/10.1016/j.rasd.2013.07.005

Wake, S.J., Izuma, K., 2017. A common neural code for social and monetary rewards in the human striatum. Soc. Cogn. Affect. Neurosci. 12, 1558–1564. https://doi.org/10.1093/scan/nsx092

Warnell, K.R., Sadikova, E., Redcay, E., 2018. Let’s chat: developmental neural bases of social motivation during real-time peer interaction. Dev. Sci. 21, 1–14. https://doi.org/10.1111/desc.12581

Wing, L., 1997. The autistic spectrum. Lancet 1761–1766.

Xiao, Y., Alkire, D., Moraczewski, D., Redcay, E., 2022. Developmental differences in brain functional connectivity during social interaction in middle childhood. Dev. Cogn. Neurosci. 54. https://doi.org/10.1016/j.dcn.2022.101079

You, X., Norr, M., Murphy, E., Kuschner, E.S., Bal, E., Gaillard, W.D., Kenworthy, L., Vaidya, C.J., 2013. Atypical modulation of distant functional connectivity by cognitive state in children with Autism Spectrum Disorders. Front. Hum. Neurosci. 7, 1–13. https://doi.org/10.3389/fnhum.2013.00482

## Reference

Esteban, O. et al. (2019) ‘fMRIPrep: a robust preprocessing pipeline for functional MRI.’, Nature methods, 16(1), pp. 111–116. doi: 10.1038/s41592-018-0235-4.

Kaufman, A. S. (2004) ‘Kaufman brief intelligence test–second edition (KBIT-2)’, Circle Pines, MN: American Guidance Service.

Lord, C. et al. (2012) Autism diagnostic observation schedule (ADOS-2). 2nd editio. Los Angeles: Western Psychological Services.

McNaughton, K. A. et al. (2023) ‘Social-interactive reward elicits similar neural response in autism and typical development and predicts future social experiences’, Developmental Cognitive Neuroscience, 59(January), p. 101197. doi: 10.1016/j.dcn.2023.101197.

